# High-throughput single-cell phenotypic profiling and backtracing exposes and predicts clinically relevant subpopulations in isogenic *Staphylococcus aureus* communities

**DOI:** 10.1101/2023.11.02.562170

**Authors:** Jonathan Hira, Bhupender Singh, Tirthankar Halder, Anel Mahmutovic, Clement Ajayi, Arif Ahmed Sekh, Kristin Hegstad, Mona Johannessen, Christian S. Lentz

**Affiliations:** Centre for New Antibacterial Strategies (CANS) and Research Group for Host-Microbe Interactions, Department of Medical Biology, UiT – The Arctic University of Norway, 9037 Tromsø, Norway; Early Biometrics & Statistical Innovation Data Science & AI AstraZeneca, Biopharmaceuticals RD AstraZeneca, Sweden; XIM University, Bhubaneshwar, India; Norwegian National Advisory Unit on Detection of Antimicrobial Resistance, Department of Microbiology and Infection Control, University Hospital of North Norway, 9038 Tromsø, Norway

## Abstract

Isogenic bacterial cell populations are phenotypically heterogenous and may include subpopulations of antibiotic tolerant or heteroresistant cells. The reversible nature of these phenotypes and lack of biomarkers to differentiate functionally different, but morphologically identical cells is a challenge for research and clinical detection. To overcome this, we present ‘<u>C</u>ellular <u>P</u>henotypic <u>P</u>rofiling and back<u>Tr</u>acing (CPPT)’, a flexible fluorescence-activated cell sorting platform, that uses optical probes to visualize and quantify cellular traits and connects the resulting phenotypic profile with a cell’s experimentally determined fate in single cell-derived growth and antibiotic susceptibility analysis. By applying CPPT on *Staphylococcus aureus* populations we recorded phenotypic signatures for dormant cells, exposed microanatomy-independent bimodal growth patterns in colony-derived cells, and revealed different culturability of single cells on solid compared to liquid media. We demonstrate that vancomycin-bodipyFL marks cellular subpopulations with increased likelihood to survive antibiotic exposure, showcasing the value of CPPT for discovery of clinically relevant biomarkers.

## Introduction

Experimental microbiology and clinical laboratory routines have traditionally studied bulk populations, revealing the behaviour of ‘saverage cells’. However, bacterial pathogens cause infections as populations comprising myriads of single cells that adapt to changing microenvironments in different biological niches or in response to antibiotics. Within these populations, cellular phenotypic heterogeneity is generated through intrinsic (e.g. stochastic gene expression, cell age) and external factors (microenvironment, cell-to-cell interactions)^1^ affecting diverse traits related to bacterial physiology and stress response^2^, antimicrobial susceptibility^3,4^, or virulence^5,6^. Heterogeneity may benefit cell populations through cooperative behaviours (division-of-labour^1,57^, sharing of extracellular resources^5,8^), as well as through generation of specialized cells with fitness advantages under adverse conditions (bet-hedging^1,5^). Alternatively, heterogeneity may result from necessary trait adjustments according to ‘cellular vigor’ ^6^. An increasing body of literature documents the clinical relevance and complications for treatment elicited by subpopulation phenotypes, particularly during chronic infections ^9,10,11-15^. This includes heteroresistant cells, i.e. cellular subpopulations in clonal isolates characterized by largely different sensitivity towards certain antibiotics^3,9,16^. It also includes non-growing, dormant cells known as persister cells that survive antibiotic treatment without developing inheritable, genetic resistance^17,18^. After antibiotic treatment, persisters can revert to a proliferating and antibiotic susceptible phenotype, can cause relapse and chronicity of infection^10,19,20^, and promote the evolution of antimicrobial resistance^21^. Another dormancy phenotype referred to as ‘viable-but non-culturable (VBNC)’ was identified by observing a discrepancy between colony forming units (CFU) and cells classified as *alive* with the help of viability stains^22,23^. The degree to which these different dormancy states are identical, related, distinct, or even artificial, is a matter of ongoing controversy^24-27^.

The presence of both dormant and heteroresistant variants is inferred retrospectively by their ability to grow under conditions where the bulk of the cells do not ^4,17^. Since there are no biomarkers to differentiate and separate these growth phenotypes from functionally different bulk cells, our ability to study them is limited. Phenotypic heterogeneity has been visualized by introduction of fluorescent labels and fluorescent reporter strains, providing increased understanding of heterogeneity particularly at the level of gene regulation^6,28-30^. In addition, we and others have shown that fluorescent chemical probes are excellent tools to study phenotypic parameters at the single cell level and expose phenotypic heterogeneity within native, isogenic cellular populations^31-33^. Coupled to time-lapse quantitative microscopy in microfluidic systems such as the mother machine, fluorescent labels and reporters are powerful tools to decipher transcriptional status and even correlate it with growth parameters ^34-38^ allowing microscopic identification and tracking of single dormant cells^37^. One disadvantage of microfluidics systems is a relatively low throughput and the difficulty to purify/recover cells for follow-up studies beyond microscopy.

To fully expose and understand cell individuality and cooperativity, a systematic framework is needed that combines visualization of diverse phenotypic traits with separation and broad functional characterization of phenotypically different cells. Inspired by pioneering studies utilizing flow cytometry-based enrichment of persisters ^39-41,24,42^, we here present a fluorescence-activated cell sorting (FACS)-based platform for high-throughput (HT) single cell phenotypic profiling coupled to functional analysis of single bacteria (**Figure 1)**. In a first stage, bacteria with different phenotypic traits are differentiated through *<u>C</u>ellular <u>P</u>henotypic <u>P</u>rofiling* (CPP) using fluorescent chemical probes, before single cells are sorted for separate analysis downstream and determination of their cell fate. In a feature we are referring to as *Phenotypic Backtracing*, the phenotypic profile of a cell at the time of sorting can be traced back after determining the cell fate post-sorting. In this proof-of-principle study focusing on the clinically relevant Gram-positive pathogen *S. aureus*, this combined platform, *<u>C</u>ellular Phenotypic Profiling and back<u>T</u>racing* (CPPT), readily detects growth variants such as non-stable small colonies and other bistable growth phenotypes by exposing the differential ability of single cells to outgrow in liquid or solid media. We demonstrate that CPP using a fluorescent vancomycin conjugate is a biomarker for reduced susceptibility to vancomycin in a vancomycin-intermediate susceptible (VISA) strain, highlighting the potential of chemical probes for differentiating clinically relevant subpopulations.

**Figure 1.**
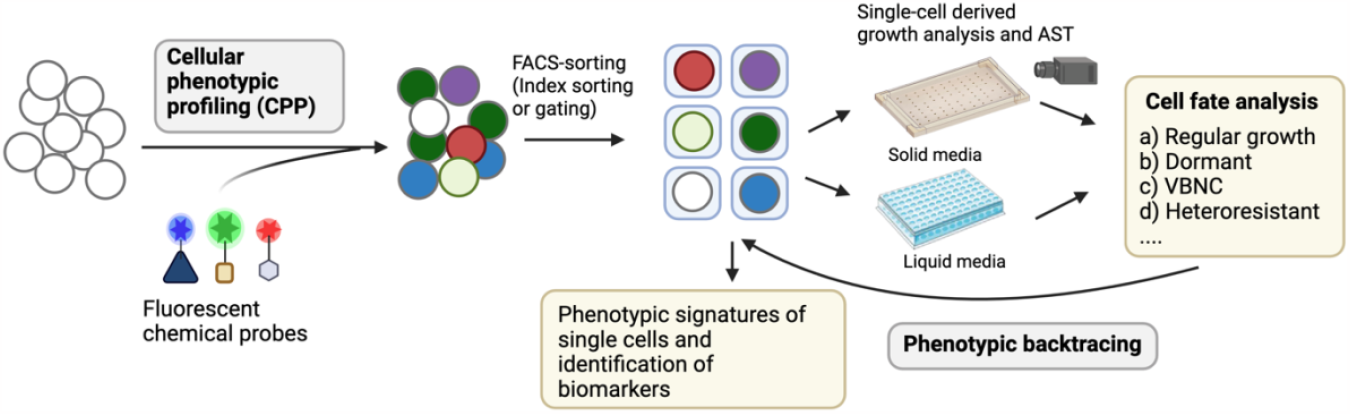
Overview of the ‘Cellular phenotypic profiling and backtracing’ (CPPT) platform. Phenotypic traits in naïve bacterial cell populations will be visualized using fluorescent chemical probes and quantified by flow cytometry. Single cells are sorted out and fed into a downstream analysis of growth performance in liquid media, where replication is monitored in a plate reader, or onto agar, where growth is monitored by time-lapse imaging. After functional analysis, cell fates are classified into categories of interest and the phenotypic profile at the time of sorting is traced back, allowing for detection of phenotypic signatures associated with dynamic reversible phenotypes and discovery of biomarkers. The figure was created with BioRender.

## Results

### Establishment of a high-throughput platform for cellular phenotypic profiling and single-cell derived growth analysis

We chose to use bacterial colonies, as they represent ideal model communities comprising bacteria differing in age, availability of nutrients, and exposure to other microenvironmental parameters that induce phenotypic heterogeneity in e.g. gene expression, replication status, or antibiotic tolerance ^43-46^. Since *S. aureus* is well known for its ability to aggregate and to form grape-like structures, we first aimed to determine whether it was possible to reliably sort single *S. aureus* cells by FACS. For method establishment, we used colonies of a plasmid-based transcriptional reporter strain producing a robust green fluorescent signal (GFP) under control of the *sar*A promoter^47^ (**Figure 2A**) and labeled cells additionally with propidium iodide (PI) to exclude dead cells prior to FACS analysis. The most commonly used flow cytometry parameter for cell size, forward scatter (FSC)^48^, failed to discriminate individual *S. aureus* cells from background buffer noise/smaller contaminating particles (**Figures S1A, 1C-iii**). However, the log side-scatter (SSC) profile reliably resolved submicron sized beads (**Figure S1A**), and *S. aureus* cells showed a narrow distribution distinct from buffer noise (**Figure S1 BC-iv**). To test if those cellular events with a higher SSC-signal might include aggregates, we sorted SSC^low^ (population P1) and SSC^high^ (P2) events onto agar for determination of colony forming units (CFU) after overnight incubation. We observed that the SSC^low^ population yielded >99%±1.4 (av. ± sd) single CFUs and the SSC^high^ population contained >89%±11.16 CFUs (**Fig. 2B, Fig. S1E,F**). Whereas the cell number can be readily determined as CFUs on solid media, in liquid culture the number of inoculated bacteria affects the lag phase.^49^ We therefore sorted single or multiple events into 96-well plates with liquid media and recorded growth curves and calculated lag phase and generation time, which for single-event derived cultures were surprisingly heterogeneous. Yet, we observed an expected inoculum-dependent decrease in lag phase (**Fig. S2**). Lag phases derived from either single or two sorted events (from the SSC^low^-P1 population) were overlapping due to the high level of heterogeneity, but statistically different (P<0.0001; **Fig. 2 B-iii**,). The lag phase from single SSC^high^-P2 events was not different to that of single SSC^low^-P1 cells, but significantly higher than that achieved from cultures derived from two SSC^low^-P1 events (P = 0.0021), suggesting that any putative cell aggregates in this SSC^high^ population either have growth characteristics indistinguishable from single CFUs or their abundance in the population is too low to affect the statistical analysis between the groups.

**Figure 2.**
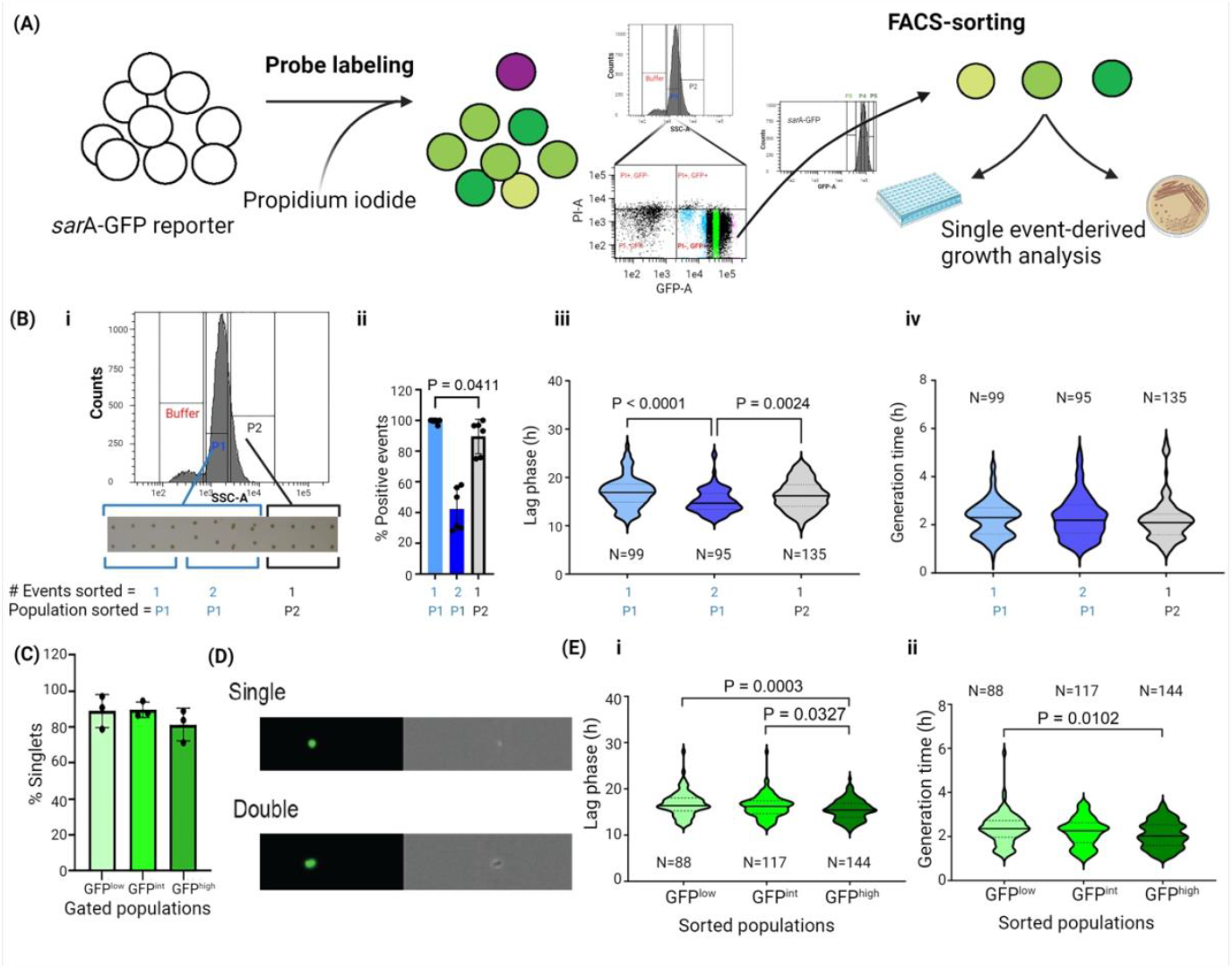
Evaluation of single bacterial cell sorting of *sar*A-GFP reporter strain. (A) Schematic overview of the experimental set up. Cells from a single *S. aureus* harboring *sar*A-GFP reporter were stained with PI and sorted into GFP^low^, GFP^int^ and GFP^high^ PI^-^subpopulations. (B) Gating *S. aureus* population for single cell sorting, (i) SSC histogram of *S. aureus* cells showing buffer noise (red) and the two cell populations SSC^low^-P1 (blue) and SSC^high^-P2 (black). The inset below shows CFUs on solid media following indicated sorting events from P1(SSC^low^) and P2 (SSC^high^), (ii) Percentage of positive events (single or double CFUs) on solid media (means ± standard deviation (SD) of 2 independent biological replicates). (iii) Violin plot showing lag phase and (iv) generation time of sorted events in liquid media. N = 95 – 135 post-sorting cultures derived from 2 independent cultures (pre-sorting). (C) Imaging flow cytometry analysis of percentage of singlets in GFP^low^, GFP^int^ and GFP^high^ subpopulations. The graph shows means ± SD of 3 independent biological replicates. (D) Exemplary photographs of singlets and doublets (GFP - left; brightfield - right) taken by ImageStream. (E) Lag phase (i) and generation time (ii) of GFP^low^, GFP^int^ and GFP^high^ subpopulations. The graph shows violin plots generated from N = 99 – 144 individual cultures derived from 3 independent cultures (pre-sorting)-Statistical significance testing in B and E, were performed by Kruskal-Wallis test in combination with Dunn’
ss multiple comparison test. N: sample size in violin plots. The figure was created with BioRender.

These data suggest that we can localize and sort single CFUs, but we cannot rule out that a CFU may be constituted by two or more cells. To address this, we separated live SSC^low^-P1 cells based on *sar*A reporter activity (GFP^+^, PI^-^) into three subpopulations, GFP^low^, GFP^int^ and GFP^high^ (**Fig. S1D**) and sorted approximately 60,000 cells per gate for imaging flow cytometry. For all subpopulations >80% of sorted events were single cells, <20% were doublets (**Fig. 2 C,D, Fig. S3**). No larger aggregates were observed. The doublets might in some cases represent the ‘smallest sortable units’ for cells in late stages of cell division or otherwise tightly interacting cells.

Although the *sar*A-dependent fluorescence reporter signal has primarily served as a simple means for detecting cells, we also wondered if our methodology might reveal biological insights into the physiological status of cells with differences in reporter signal. We found that growth of GFP^low^ cells was significantly delayed compared to cultures derived from GFP^int^ or GFP^high^ cells (**Fig.2 E**). Of note, cultures derived from GFP^low^ cells also demonstrated more heterogenous growth kinetics and a shift towards higher generation times. These data suggest that cells with higher *sar*A transcriptional activity replicate more rapidly, but it must be considered that the results may be confounded by plasmid copy number that may affect both reporter signal and growth kinetics on antibiotic selection media. Nevertheless, our proof-of-concept data reveal that the phenotypic state of the cell at the time of sorting indeed translates to different growth-related characteristics read out hours later.

### Cellular phenotypic profiling coupled to single cell-derived growth analysis detects bimodal growth phenotypes independent of colony microanatomy

To test if our platform can detect growth heterogeneity without the confounding factors associated with the plasmid-based reporter strain, we decided to study wildtype (WT) cells from single colonies of bacteria of the MRSA strain USA300 LAC. Cells were phenotypically profiled using PI and the metabolic reporter dye Redox Sensor Green™ (RSG) (**Fig. 3A**), which is activated by reductases in the respiratory chain (**Fig. 3A**). In contrast to the sharp fluorescence signal achieved with the *sar*A-GFP reporter strain, the RSG signal showed a wide distribution that was dose-dependent (**Fig. 3A, Fig. S4**) and exposed two distinct probe-labelled subpopulations (RSG^low^, RSG^high^) and a minor RSG^(-)^ population. Cells from these subpopulations were sorted for single event-derived growth analysis in liquid media. The first notable observation was that RSG^(-)^ events led to growth suggesting they are live cells, despite their inability to stain with the metabolic dye. One possible explanation could be limited permeability of the probe in this population. Next, when comparing the growth kinetics of different single-cell derived cultures, we observed large cell-to cell heterogeneity in lag phase and a bimodal distribution with peaks at ≈ 10 h and 14 h, respectively (**Fig. 3B**). In contrast, generation time was highly similar for all single cell-derived cultures (**Fig. 3C**). Our data thus suggest the presence of two physiologically distinct subpopulations in the cell population pre-sorting. Since cultures derived from RSG^(-)^, RSG^low^ and RSG^high^ cells all showed similar bimodal distribution patterns, we conclude that the downstream growth performance is independent of the metabolic/redox-status reported by RSG during sorting (**Fig. 3 B, C**).

**Figure 3.**
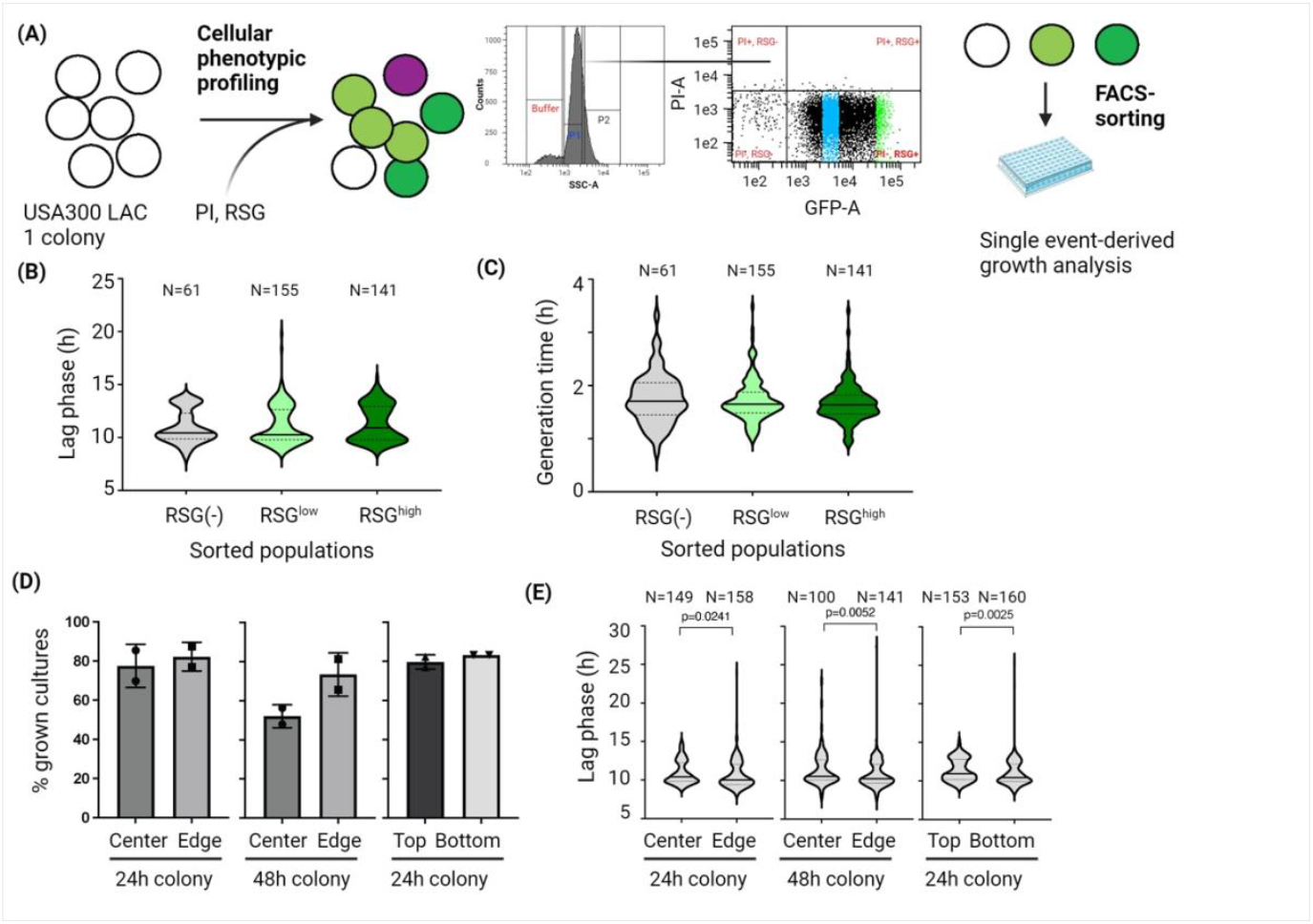
Heterogeneity of growth phenotypes in *S. aureus*. (A) Schematic overview of the experimental setup. Cells from a single USA300 LAC colony were stained with RSG and PI. RSG^(-)^, RSG^low^ and RSG^high^ subpopulations were sorted for single-event derived growth analysis. (B-C) RSG phenotype pooled from 3 different biological replicates (pre-sorting): Lag phase (B) and generation time (C) of single-cell derived cultures derived from RSG^(-)^, RSG^low^ and RSG^high^ subpopulations. No significant differences between the groups based on Kruskal-Wallis testing in combination with Dunn’s multiple comparisons test. (D-F) Single-cell derived growth performance of unlabelled LAC cells harvested from different locations of a colony. (D, E) Bacteria were harvested from either center and edge or top and bottom of a colony grown for 24 or 48h and subjected to single-cell derived growth analysis in liquid culture. (D). Percentage of positive cultures 48h growth post single event sorting. The bars represent means (± SD) of two independent experiments (indicated data points) that were derived from 96 sorted events each. (E) Violin plot showing lag phase of single-event derived growth cultures pooled from two independent experiments. Cross line: Median lag time. Statistical differences between the groups were calculated pairwise by two-tailed, non-parametric Mann-Whitney test. The figure was created with BioRender.

We wondered if this bimodality could be explained by the microanatomy of the colony, *i*.*e* that cells in the center vs. edge might differ in cell age and lag phase, or whether direct nutrient availability influences growth kinetics of cells in the top or bottom part of the colony. We therefore harvested cells from different locations and compared their lag phase and growth kinetics. We found that cultures harvested from the center of the colony included a higher percentage of events that did not result in downstream growth post sorting, suggesting the presence of either dead or VBNC cells (**Fig. 3D**). Cells from the bottom and edge had lower lag phases than cells from the centre or top, respectively (**Fig. 3E**). However, the bimodality was observed throughout all four conditions, and the location of the cells in the colony did not influence their outgrowth kinetic, suggesting that this level of heterogeneity is not primarily dictated by the location and thus microenvironment of the cells.

### *Phenotypic backtracing* connects identification of dormant subpopulations with single-cell phenotypic profiles

Having dissected heterogenous growth phenotypes in colonies, we sought to address dormant, i.e. non-stable small colony phenotypes that can be elicited by prolonged growth at low pH ^42 13^ (**Fig 4A**). Under these conditions we found an increase in putatively dead, PI^+^-cells (11.6 ± 0.8%, n=3), while the two subpopulations of RSG^low^ and RSG^high^ cells were still evident. Monitoring colony formation for 48 h revealed that 66.5 % of sorted events gave rise to ‘regular’ colonies, whereas 6.25 % of events gave rise to small/delayed colonies. In other words, 8.6% of all formed colonies were small which concurs with previous observations by Huemer et al. ^13^. In 27.25 ± 5% of sorted events no colony appeared within 48 h (**Fig. 4B, S5Bi**). These cells might be in a deeper state of dormancy or VBNC-state ^50,51^. Interestingly, we observed individual colonies that appeared as late as 72 h after sorting (**Figure S6**), suggesting that the population of events that does not give rise to colonies by 48 h indeed contains viable cells that become ‘culturable’ later.

**Figure 4.**
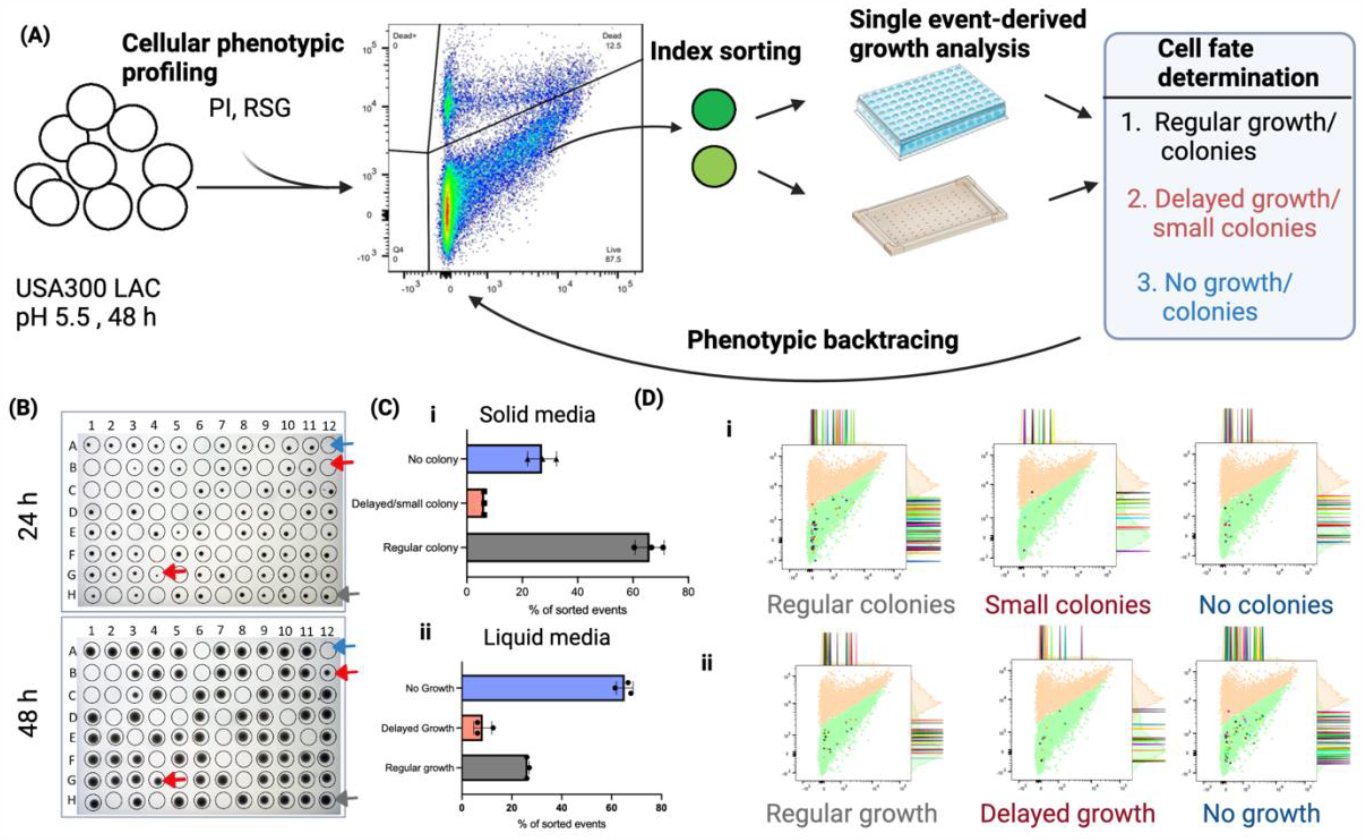
Phenotypic signatures of *S. aureus* subpopulations: (a) Schematic overview illustrating the experimental setup. After cultivation in media with low pH for 48 h, *S. aureus* USA300 LAC cells were labeled with RSG (for live cells) and PI (for dead cells), followed by flow cytomety analysis. Using an index sorting approach that enables phenotypic backtracing, live cells (PI^-^) were subsequently sorted for single-cell derived growth analysis on agar and in liquid media. (b) Exemplary photographs of individual events sorted as single cells on a single-well agar plate. The top image shows colony growth after 24 hours, while the bottom image displays colony growth after 48 hours. Different cell fates are highlighted with colored arrowheads: Normal (gray), delayed/small colonies (red), and no growth (blue). (c) Bar graph demonstrating distinguishable growth phenotype features on solid media (i) and liquid media (ii). Bars show means ± s.d. of n=3 biological replicats. Dots: individual replicates. (d) Indexed dot plot generated using the index sort package from Flowjo software. This dot plot reveals the ancestral locations of individually sorted events based on their phenotypic signatures, as determined by RSG for both growth conditions on i) solid and ii) in liquid media. The figure was created with BioRender.

Dormant growth variants have commonly been observed on solid media but are impossible to detect with traditional bulk-based liquid culture techniques. Our experimental platform, in contrast, allows for single cell-derived outgrowth studies in liquid culture and revealed that the percentage of sorted events that were not able to yield detectable growth was more than twice as high (65 ± 5%) compared to events that failed to grow on solid media (**Figure 4B, C**). Thus, not all cells that give rise to colonies on agar are able to grow in liquid media.

One common limitation in the field of persisters/non-stable small colonies is that their phenotype can only be inferred retrospectively once cells have reverted from dormancy into a proliferating state. To overcome this limitation, we developed *phenotypic backtracing* (**Fig. 4A**). Phenotypic backtracing connects the fate of a cell post-sorting (e.g. after determining whether the cell was dormant or not) with a phenotypic characterization of a cell pre-sorting (*e*.*g*. while a cell *is* dormant). Here, we classified the growth phenotypes displayed by cells post-sorting into three cell fate categories: i) regular growth, ii) delayed growth, iii) no growth. For each sorted event we could *trace back* the founder cell in the FACS-plot, thus connecting a cell’s fate with its phenotypic profile (**Fig. 4D**). We observed that cells from all cell fate categories showed a similar distribution in the RSG vs. PI profile. Thus, the bimodal RSG signal is not correlated with dormancy status of *S. aureus* under these conditions. This finding concurs with previous observations that although *S. aureus* persisters are associated with low ATP levels and reduced metabolic activity^41,52^, metabolism and transcription are not completely abolished^13,42^.

### Phenotypic profiling with a fluorescent antibiotic conjugate denotes isogenic subpopulations with reduced vancomycin susceptibility in vancomycin-intermediate susceptible *S. aureus* (VISA)

Whereas chemical probes for dormant phenotypes remain to be developed, some probes whose labeling profiles may correlate with the susceptibility to certain antibiotics exist. Zhang et al. demonstrated that cellular labeling profiles of *S. aureus* strains with fluorescent analogs of the glycopeptide antibiotic vancomycin were dependent on molecular characteristics of vancomycin resistance^53^. Vancomycin targets D-Ala-D-Ala in peptidoglycan precursors peptides. Vancomycin-resistant *S. aureus* (VRSA) strains produce a modified version the peptidoglycan precursor (D-Ala-D-Lac) which binds vancomycin with reduced affinity leading to reduced cellular labeling with fluorescent vancomycin analogs in VRSA compared to susceptible strains (VSSA) ^53^. In contrast, VISA strains, for which resistance is associated with decreased autolysis, increased cell wall thickening and putatively production of decoy targets ^54-57^, revealed increased labeling of vancomycin probes compared to VSSA^53^. Since VISA and heteroresistant VISA (hVISA) commonly display heterogeneous resistance patterns^58,59^, we asked if fluorescent vancomycin probes could be used in the CPPT platform to expose subpopulations with different susceptibility to vancomycin^58^ (**Fig. 5A**).

**Figure 5.**
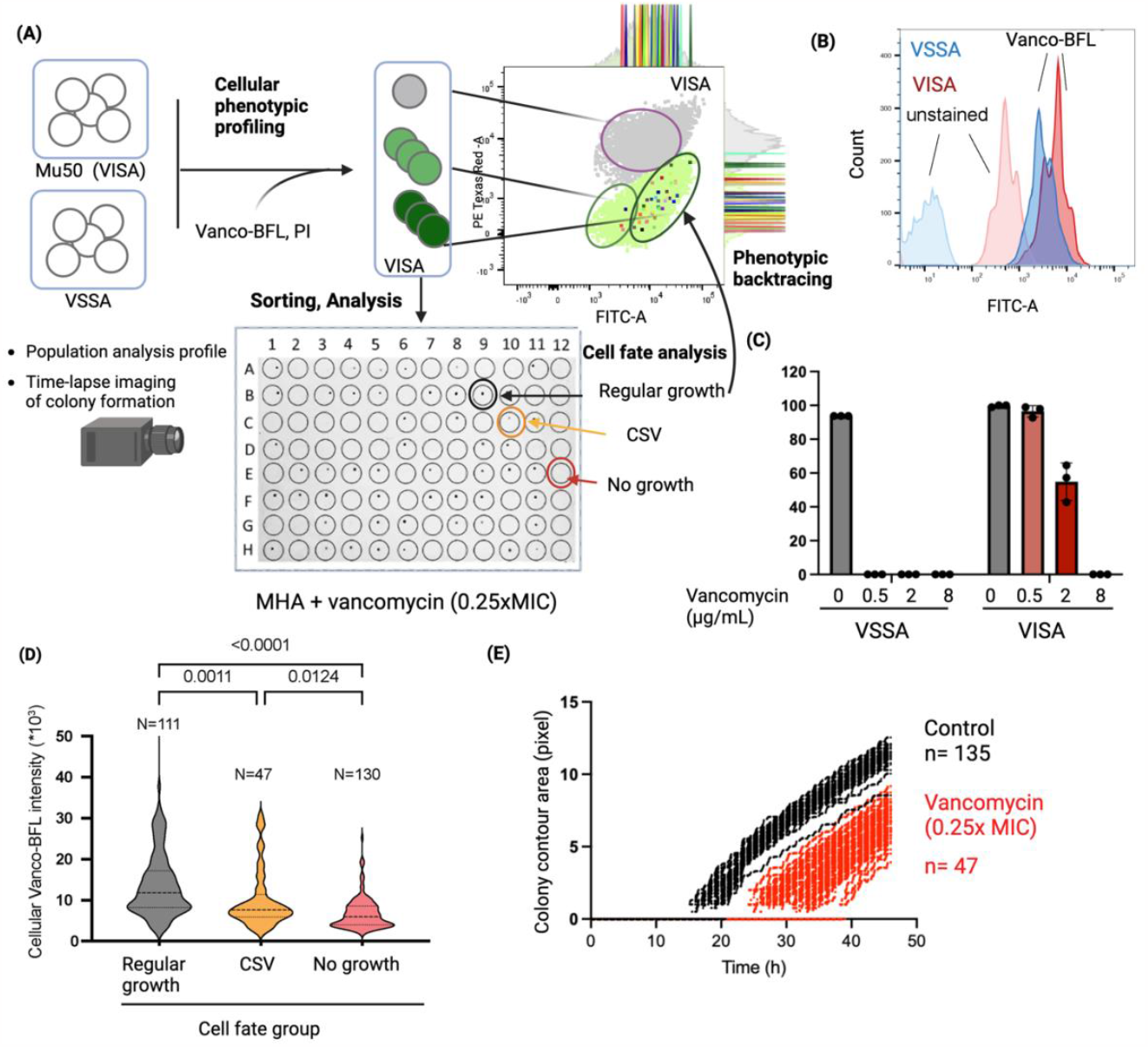
Heterogeneity in vancomycin resistance in a VISA strain. A) Schematic overview of the CPPT setup for comparative analysis of VISA strain Mu50 and a VSSA control. Cells were labelled with Vancomycin-BFL and PI. PI^(+)^-cells are depicted in gray, PI^(-)^-cells with different degrees of Vanco-BFL-labeling are indicated in light and dark green. PI^(-)^-cells were sorted onto Mueller-Hinton Agar (MHA) supplemented with different concentrations of vancomycin and colony formation was observed at different time points and for individual plates by time-lapse imaging. For VISA cells growth on a 0.25x MIC plate was used for cell fate determination (growth, no growth, and colony size variant (CSV), i.e. smaller colonies) prior to phenotypic backtracing of the ancestral phenotypic signatures in the flow cytometry dot plot. B) Flow cytometry histogram showing the fluorescence signal in the FITC-A channel of unstained and Vanco-BFL-labelled VSSA (blue) and VISA cells (red) ’. C) Percentage of sorted events that gave rise to colonies on agar plates with different concentrations of vancomycin. (D) Violin plot showing the cellular Vanco-BFL signal intensity of VISA cells for the different cell fate groups determined after growth on 0.25xMIC vancomycin. The graph shows pooled events derived from 2 independent cultures (pre-sorting) with 96 sorted events each. Statistical significance testing was performed by Kruskal Wallis test in combination with Dunn’s multiple comparisons test. (E) Colony growth dynamics of sorted VISA cells in the presence (0.25xMIC) and absence of vancomycin as analyzed by time-lapse imaging analysis. Growth (Y-axis) was measured as contour size on the edge of the colony and is plotted against cultivation time. The figure was created with BioRender.

We compared the labeling profiles of the VISA strain Mu50^60^ and a VSSA control with the commercially available fluorescent probe Vancomycin-BODIPY FL (Vanco-BFL). Consistent with the observation by Zhang et al. ^53^, we found that VISA cells labelled more strongly (**Fig. 5B**). Interestingly, also in the absence of probe, VISA cells showed increased autofluorescence in the fluorescein channel, which may be attributed to its increased cell wall thickness ^55,56^. The flow cytometry profile further indicated subpopulations with different Vanco-BFL signal in the VISA cell population. To determine if subpopulations of the VISA strain differ in their antibiotic susceptibility we sorted single cells onto agar with different concentrations of vancomycin, thereby establishing a traceable variation of the population analysis profile (PAP) test which is commonly used to detect heteroresistance ^57^. Whereas cells grew uniformly at 0.0625xMIC vancomycin and no colonies formed at 1xMIC (=8 μg/mL), at 0.25xMIC 55% of the single cells grew to colonies supporting the existence of subpopulations with different levels of susceptibility to vancomycin (**Fig. 5A,C, Fig. S7**). To test if this observed heterogeneity in vancomycin-susceptibility was correlated with the cellular Vanco-BFL labeling profile, we implemented *phenotypic backtracing*. Sorted events were assigned into three cell fate categories depending on their ability to grow and the size of the resulting colony. Although all VISA colonies recovered on 0.25x MIC were generally smaller compared to colonies grown in the absence of vancomycin, we are referring to the bulk of these colonies as ‘regular growth’, and to a subpopulation of colonies with further reduced size as ‘colony size variants’. Phenotypic backtracing analysis revealed that cells displaying regular growth localized to a specific area of the Vanco-BFL– PI plot and showed significantly higher f Vanco-BFL signals compared to cells from the other cell fate groups (**Fig. 5A,D; Fig. S8A**). We conclude that Vanco-BFL has utility as a biomarker that denotes a subpopulation of cells that show an increased likelihood of displaying phenotypic resistance. Of note, increased labeling alone is not sufficient to explain this resistance since a fraction of cells with high levels of Vanco-BFL labeling was still unable to grow (**Fig. 5B**). Since colonies recovered in the presence of 0.25xMIC vancomycin had a reduced size, we implemented an in-house built solution for automated time-lapse imaging of colony formation post-sorting, adapted from ColTapp^61^, to determine if a) surviving cells were sufficiently pre-adapted to grow in the presence of vancomycin but do so at a slower rate (explaining the small colony size) or b) if cells needed to undergo additional steps of functional adaptation, leading to a delayed onset of growth. The experiment revealed that in the presence of vancomycin colonies became detectable on average 12-14 h later than in the absence of antibiotic (**Fig. 5E, Fig. S8B**), but once detectable, colonies grew at similar rates in the presence and absence of vancomycin. Collectively, our results indicate that the cellular trait causing increased labeling with Vanco-BFL is a necessary, but not sufficient precondition to survive antibiotic exposure, and that the ability to grow in the presence of vancomycin required additional functional adaptations inducing a growth delay. This finding concurs with reports on changes in metabolism and cell wall biosynthesis gene expression in VISA strains upon vancomycin exposure^62^. It remains to be determined if differences in cell wall structure, such as absence of glutamine amidation and increased cell wall thickness which were reported as general characteristics of Mu50 ^56^ are also the molecular basis for the observed heterogeneity in Vanco-BFL labeling and vancomycin-susceptibility.

## Discussion

Understanding cell individuality is an important precondition to understand cooperative behavior in bacterial populations. When individual cells are morphologically identical, which is the default scenario for most isogenic populations, differentiating functionally different cell is a formidable challenge. In recent years several technical advances have been made. Single-cell sequencing approaches, for example, provide in-depth and high-throughput (HT) information about the transcriptional status of cells^63,64^, but their use requires the destruction of cells, preventing further functional analysis. On the other hand, microfluidic studies have emerged as the gold standard for visualizing specific phenotypic traits using fluorescent markers while allowing for a microscopy-based assessment of replication and antibiotic susceptibility in real-time ^34,35,38^. Yet, the general throughput is limited, functional analysis largely restricted to microscopy and analysis of bacteria whose growth requires cell division events in multiple planes is challenging. We are convinced that our flow cytometry-based CPPT-platform provides valuable, complementary information for deconstructing cellular phenotypic heterogeneity in bacterial populations. Whereas microfluidics studies directly track single cells and their replication microscopically, CPPT differentiates and separates single-cells, but the growth kinetics are detected in more traditional growth assays with the distinction that they are single cell-derived. In the future, this observation gap may be closed by combining cell sorting with downstream real-time-microscopy in microfluidic systems.

In combination with simple single-cell derived growth assays, CPPT has exposed previously unidentified heterogeneity-related phenomena, such as the bimodal distribution of lag phases not related to colony topography, or the differential ability of dormant single cells to outgrow on agar, but not in liquid culture. The latter finding represents a new nuance in bacterial cultivability and adds another level of complexity and additional fuel to the ongoing debate of when cells are viable, culturable, dormant, or dead.^24-27^ Our data also suggest that caution must be taken when comparing studies addressing dormancy and cultivability in agar-based systems, compared to e.g. microfluidics-based studies performed in a (confined) liquid environment. While the mechanisms underlying these newly described phenotypic variants remain to be determined, these results illustrate the power of CPPT to uncover fundamental new aspects of bacterial physiology and ecology. Our flow cytometry-based platform is tunable to operate at the single-cell level, as in this work, but can also be used with gating to sort subpopulations with a specific labeling profile for diverse downstream assays, such as ‘omics’ studies or infection or virulence-related assays.

CPPT also provides new opportunities for dissecting known cellular subpopulations. For persister research, one experimental dilemma is that the phenotype can only be assigned retrospectively, after cells have undergone a phenotypic reversal to a replicating state. Our platform provides a potential solution that combines determination of the cell fate (e.g., to confirm dormancy by the ability of cells to grow after an increased lag), with backtracing to cellular phenotypic profiles determined by flow cytometry. This strategy allows for broad-scale phenotypic characterization of dormant cells using diverse combinations of fluorescent chemical probes that could enable identification of markers for the direct detection and enrichment of dormant cells. The feasibility of this strategy is supported by our finding that Vanco-BFL can differentiate clinically relevant subpopulation phenotypes and marks a subpopulation of cells with increased vancomycin-resistance. This clear distinction is remarkable since strain Mu50 is considered more homogenous in its resistance profile than hVISA strains^65^, for which even more pronounced differences in labeling may be expected.

Chemical Biology is providing an ever increasing toolset of next-generation physiology probes that report on diverse phenotypic traits from cellular permeability, specific biomolecular uptake pathways, replication, metabolic activity or the distribution and activity of specific molecular targets or groups in living bacterial cells (as reviewed in ^33,66,67^) and are exploitable for CPP. The use of exogenous fluorescent probes makes CPP highly compatible for applications in clinical samples^15^, allowing for direct detection and characterization of unstable phenotypic variants that would revert upon cultivation. We believe persisters and heteroresistant cells are only the most easily detected tip-of-the-iceberg of distinct subpopulation phenotypes in isogenic bacterial pathogen populations. A systematic implementation of CPPT will help uncover further clinically relevant subpopulation phenotypes, understand their ecological role and interplay with other cells in the populations, and enable development of strategies to detect, target, or manipulate subpopulation phenotypes to improve the clinical outcome of bacterial infection management and antimicrobial treatment.

## Materials and Methods

### Bacterial strains and Growth conditions

*Staphylococcus aureus* strains analyzed in this work are summarized in **Table 1**. Wildtype strains were routinely cultivated in Tryptic Soy Broth/Agar at 37°C unless stated otherwise. The fluorescent *sar*A reporter strain transformed with pCM29-p*sarA*-GFP (courtesy of Dr. Alexander Horswill)^47^ was grown on Tryptic Soy Agar plates supplemented with 10 μg/ml chloramphenicol.

**Table 1.**
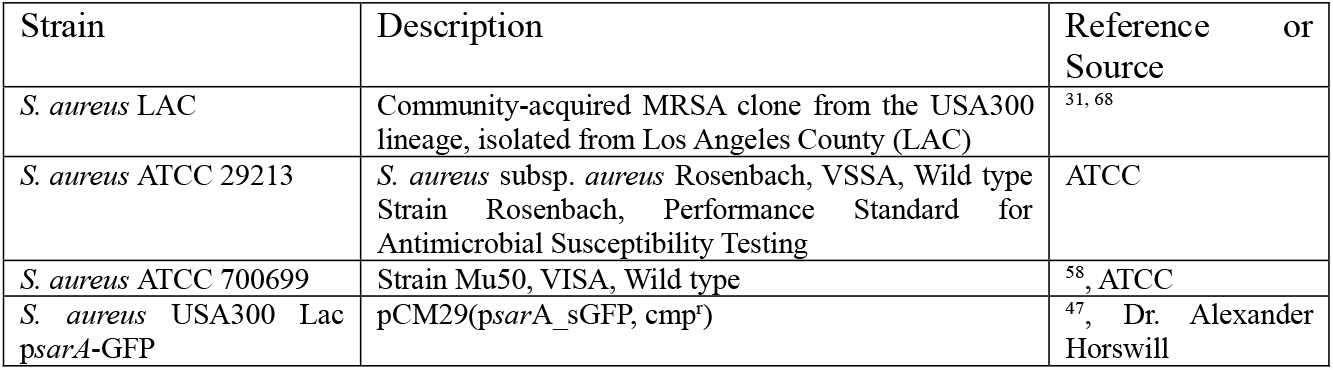
Strains used in this study.

### General flow cytometry and cell sorting

All samples in this research were analyzed by flow cytometry and sorting instrument BD FACSAria III instrument (BD Biosciences). The instrument was configured and pre-calibrated with BD® CS&T beads according to the manufacturer’s protocols to conduct quality assessments of the instrument’s optical, electronic, and fluidic systems, as well as facilitate the calibration of fluorescence compensation. Before sorting, BD FACS™ Accudrop Beads were used to ensure proper drop formation and sort accuracy during the flow cytometry and sorting process. A neutral density (ND) filter 1.0 was used to regulate the laser power and optimize signal-to-noise ratio during data acquisition. Importantly, 70 μm nozzle with a sheath pressure of 70 psi was used for sorting to achieve optimal separation of cells and minimize clogging of the instrument. All other parameters such as FSC (Forward scatter), SSC (Side scatter), fluorescence channels and their voltage, threshold were adjusted as needed based on experiments and optimization. Flow rates and sort purity during sorting were maintained at a consistent and optimal level to ensure efficient and accurate separation of cells and adjusted based on sample and sorting objective. Both Instrument setting and data acquisition was performed using BD FACSDiva software V 9.0 and FlowJo software (V 10.8.1) were used for data analysis.

### Imaging flow cytometry

Imaging flow cytometry was performed on an ImageStreamX MkII (Amnis). The detailed procedures are described under specific experimental procedures for proof-of-principle studies with fluorescent reporter strain below.

### Phenotypic backtracing

We employed the phenotypic backtracing method to analyze the observed ancestral cell functions that translates into functional traits exhibited by bacterial colonies at single cell resolution. In this pipeline, backtracing of cell function was implemented through indexing of individual cell, which are to be sorted. Index sorting represents a FACS sorting mode that enables the isolation of individual bacterial cells, while allowing for a retrospective assessment of all fluorescence and scatter parameters associated with each cell. BD FACSAria III instrument was used for index sorting. The indexing parameter were set to single cell precision with target event 1 and the sorting layout to 96 well plate. Single cell precision does not allow interrogated drop to be sorted if 2 target cells are present. Index data was further analyzed with FlowJo software (V 10.8.1) and its indexSort package. This package retrieve index sorted data from fcs data files and visually explore for downstream analysis.

### Single-or multiple event derived growth kinetics in Broth media

Single or multiple flow cytometry events (i.e. cells) were directly sorted into individual wells of a 96-well plate (353072, Falcon) containing 200 μl of either TSB (WT) or TSB supplemented with chloramphenicol (for *sar*A-GFP reporter strain) based on experimental condition. Plates were then sealed with Breathe-Easy sealing membrane (Z380059, Merck) and incubated in the Biospa 8 (Biotek) at 37°C and 80% humidity. Absorbance at 600 nm (A600) was measured every 30 minutes for a period of 48 h using Synergy H1 (Biotek) plate reader. Prior to each read, plates were shaken orbitally at low speed for 10 s following by a 5 s pause. Biospa 8 and Synergy H1 were operated via Biospa on Demand and Gen 5.3.10 software, respectively. Data were exported to an excel file (.xlsx) using Gen 5.3.10 software. Data files were curated such that they contained time in 00:00:00 format (column A), temperature (column B) and A600 values from wells A1 to H12 in subsequent columns. Headers and all other information were removed from curated .xlsx files, which were saved with an extension of _A.xlsx (e.g., FILE NAME_A.xlsx). An example of curated file is presented in supplementary files. Exported data was further analyzed using MATLAB script (see code availability), which generates growth rate (h^-1^), lag phase duration (h) and onset of stationary phase (h) and generation time (h) was calculated as LN (2)/growth rate. Data were plotted using GraphPad Prism 9.5.

### Modeling the lag phase, the stationary phase and the exponential growth rate of a microbial population of cells

We use a simple non-linear regression approach to fit a Hill equation to the log-transformed fluorescence measurements to determine the duration of the lag-time t_L_, the onset of stationary phase t_S_ and the exponential growth rate r (**Figure S2A**). Specifically, the regression equation f(t) is of the form

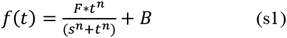

where F is the maximum theoretical fluorescence measurement relative to the baseline and therefore F+B is the absolute maximum fluorescence value. The elapsed time is labeled by t, n is the Hill coefficient that determines the sigmoidicity of the regression function and s is the time point at which the fluorescence level is at half its theoretical maximum level shifted by the baseline (**Table 2**). We define both t_L_ and t_S_ relative to the region of exponential growth, i.e. if the bacteria are not growing exponentially then they are either in lag phase or in stationary phase. The exponential growth region is defined by the intersection of the tangent line of f(t) with steepest slope with the baseline B and with the absolute maximum fluorescence value F+B. By definition, the time point at which the tangent line is steepest is given at the inflection point t_I_ (eq. s2).

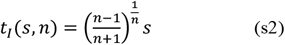

Hence the growth rate r is given by the derivative of f(t) evaluated at t=t_I_,

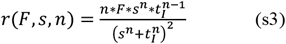

The duration of the lag time, t_L_, is given by the intersection of the tangent line at t_I_ and the baseline B given by *f*(*t*_*I*_) + *f*′(*t*_*I*_)(*t*_*L*_ − *t*_*I*_) = *B*. Solving for t_L_ with f’(t_I_)=r (eq. s2) we get

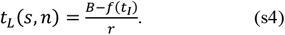

We take the same approach as we did for the lag phase to determine the time point of onset of stationary phase, t_S_, and look for where the tangent line at t_I_ intersects the absolute maximum fluorescence value (F+B). That is, we solve for t_S_ from *f*(*t*_*I*_) + *f*^′^(*t*_*I*_)(*t*_*S*_ *− t*_*I*_) = *F* + *B* resulting in

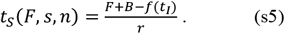

**Table 2.** Definition, meaning and bounds of variables.

| Variable | Meaning | Bounds |
| --- | --- | --- |
| **F** | Maximum fluorescence relative to baseline. | $F > 0$ |
| **B** | Baseline fluorescence level. | $B \geq 0$ |
| **n** | Hill coefficient. | $n > 1$ |
| **s** | The time point at which the fluorescence level is at $F/2 + B$. | $s > 0$ |
| **t<sub>L</sub>** | Duration of the lag phase. | $t_L > 0$ |
| **t<sub>I</sub>** | The inflection point. | $t_I > 0$ |
| **t<sub>S</sub>** | The time point at which stationary phase begins. | $t_S > 0$ |
| **r** | Exponential growth rate. | $r > 0$ |

### Growth kinetics on Agar media

Single or multiple flow cytometry events were directly sorted in a single well plate (internal dimensions 118.63*82.13mm; 734-2977, VWR) containing ∼45 ml of either TSA (for WT) or TSA supplemented with chloramphenicol (for *sar*A-GFP reporter strain), and Mueller-Hinton (MH) agar with or without antibiotic (Vancomycin). Single well plates were sorted using single cell precision mode by the adjusted 96-well sorting layouts in FACSDiva software. Following sorting, plates were placed inside of in house built real time colony tracking imaging platform. The platform comprises metal box with preinstalled cannon eos M200 camera with central temperature control condition, which at 37°C. Images from agar plate were acquired every 5 min interval for 24h (total of 289 frames) or 48hrs (578 frames) and converted into time-lapse single video file.

### Time-lapse imaging of colony growth

The resulting video files of single cell growth on agar plates were analyzed with in house-built software for tracking colony growth dynamics in real-time. The process of colony detection involves a series of image analysis operations, starting with a grayscale image. Local adaptive thresholding is applied to create a binary image, followed by artifact removal and the elimination of unwanted objects based on size criteria. Overlapping colonies are separated using watershed segmentation. Subsequently, this code employs circular filtering and contrast enhancement on isolated object images. To address false positive detections, sequential quality control functions and Mahalanobis Distance^69^ estimation are used to exclude them from further analysis. Overall, the process includes multiple steps, such as thresholding, artifact removal, watershed segmentation, object extraction, contrast enhancement, and quality control checks, ensuring accurate identification and analysis of colonies while minimizing false positives. Additionally, the base method which we developed, is employed for object tracking, which involves object detection, state propagation across frames, association with existing objects, and object lifespan management, enabling accurate tracking and analysis over time. The object model used for tracking targets across frames consists of two main components: the representation of the target and the motion model used for predicting its position in the next frame. To estimate the target’s movement, a linear constant velocity model is employed, assuming that the inter-frame displacements remain consistent over time. This model operates independently of other objects and camera motion, ensuring accurate and reliable tracking of individual targets throughout the video sequence. In the target tracking process, when a detection is successfully associated with a target, the target’s state is updated by incorporating the information from the detected bounding box. This update involves optimizing the velocity components using a Kalman filter ^70^ framework to achieve optimal estimation. However, if no detection is associated with the target in a particular frame, the target’s state is simply predicted without any correction using the linear velocity model. This hybrid approach enables accurate and continuous tracking of targets by updating their states whenever new detections are available, while also providing predictions when no new detections are found. The proposed method for tracking individual structures involves segmenting the structures and then performing Kalman filtering in image space, along with the Hungarian algorithm ^71^ algorithm using bounding box overlap as an association metric, 1 to track and estimate their sizes over time. This integrated approach allows for accurate and reliable tracking of structures throughout the temporal sequence of images, providing valuable insights into their behavior and characteristics. Estimated size of colonies was converted to pixel unit and performed fitting time-resolved data analysis using an open-source R package QurvE ^72^.

### Code Availability

All analysis performed with in house developed scripts for broth culture and time lapse agar kinetics are available at github.com.

Code for ‘single cell-derived growth analysis’ and ‘time-lapse imaging of colony growth ’ were implemented in MATLAB (R2021a), The MathWorks, Natick, MA, USA) and are available on GitHub:

https://github.com/HMIUiTL9/Colony-growth-time-lapse-Imaging.git

https://github.com/HMIUiTL9/Single-cell-derived-growth-curve.git

All scripts are free software and are free to redistribute and/or modify under the terms of the GNU General Public License as published by the Free Software Foundation, either version 3 of the License, or any later version.

### Statistical analysis

Statistical analysis was performed using Prism 9.5 or 10.0 (GraphPad software, San Diego, US). For experiments with 3 or more groups, data were analyzed by Kruskal Wallis test with Dunn’s multiple comparisons test. Pairwise analysis in experiments performed with two experimental groups were done using non-parametric, two-tailed Mann-Whitney test or parametric, two-tailed Student’s t-test, as appropriate.

### Specific experimental procedures for proof-of-principle studies with fluorescent reporter strain

#### Sample preparation

A fresh single colony of the *sar*A-GFP reporter strain grown overnight was suspended in PBS (D8537, Sigma) and mixed well using a vortexer. Suspended bacteria were sonicated for 5 minutes at room temperature (RT) using Branson 3510 water sonicator to break bacterial aggregates. The cells were diluted to 1*10e^7^ cell suspension in 1 ml PBS, were stained with 0.067 mM propidium iodide (PI, B34954, Invitrogen) for 5 minutes at RT, and were sonicated for 1 minute prior to flow cytometric analysis. Unstained cells were prepared analogously and were used to set up flow cytometer gates.

#### Flow cytometry and FACS-sorting

The blue laser (488 nm) was used to collect Forward scatter (FSC) and Side scatter (SCC) through photodiode, and GFP signals through 530/10 bandpass filter; The yellow-green laser (561 nm) was used to collect PI signals through 582/15 filter. Threshold operator or triggering threshold was set to 200 on SSC. PMT voltages were adjusted for optimal separation of different populations as 650 (FSC), 420 (SSC), 650 (GFP), and 850 (PI). The general gating strategy used in this is presented in Supplementary figure S1. Unstained cells were used to set up the gates for both GFP- and PI-positive and negative populations, respectively. First, 0.58, 0.79 and 1.3 μm beads (NPPS-4K, Spherotech) were used to adjust log intensities of side scatter (SSC) vs forward scatter (FSC) scales such that beads can be clearly separated from the buffer noise. The integrated value of area (A) is usually sufficient to resolve larger cell types such as immune cells, however, we used integrated value of height (H) for SSC and FSC which fits all events well within the plot area while clearly separating the buffer noise. SSC and FSC voltages (PMT) were adjusted for unstained cells as beads and cells of same sizes have different refractive indices ^73^. The relationship between size beads and bacterial cell population is shown in Supplementary Figure S1A,B. Beads of less than 1 μm in size showed a broad FSC profile despite uniform in size and events of 0.59 and 0.79 μm beads were difficult to distinguish. FSC profile of the bacterial showed size distribution in this range. However, the SSC-A profile showed a clear distinction between beads of different sizes. Using a conventional flow cytometer it is challenging to resolve submicron particles ^74^ and SSC has been used to clearly resolve submicron particles ^48^. SSC-A distribution of bacterial cells showed a narrow distribution which was clearly distinct from buffer noise. The P1 population (SSC^low^) was gated on SSC-A histogram around the peak area and P2 population was set to capture events with the highest SSC-A signals. Relationhips between different peaks in different plots is shown in Supplementary figure (S1C).

The P1 population was discriminated for the presence or absence of GFP by gating GFP+ or RSG+ population and for PI signal to eliminate dead cells (PI positive cells) on a dot plot of log PI-A and GFP-A intensities. Dot plot was split into four quadrants, as, Q1 (PI+, GFP-/RSG-), Q2 (PI+, GFP+/RSG+), Q3 (PI-, GFP-/RSG-), and Q4 (PI-, GFP+/RSG+). GFP/RSG fluorescence of the 4^th^ quadrant (Q4) population was displayed on a histogram of counts and log GFP/RSG-A intensity. On this histogram, low, intermediate, and high GFP intensity populations (GFP/RSG^low^, GFP/RSG^int^, and GFP/RSG^high^, respectively) were gated as P4, P5 and/or P6.

#### Downstream analysis

Single-or multiple cells were sorted onto agar plates for CFU analysis or into 96-well plates for growth analysis in liquid culture, following the general procedures described above. Fluorescently labelled bacteria from the P4, P5 and/or P6 gates were also subjected to imaging flow cytometry. At least 60,000 events per gate were sorted using FACSAria and were used as input for imaging flow cytometry (ImageStreamX MkII, Amnis). Samples were run at low speed and high sensitivity setting; images were collected using 60x magnification. We collected GFP-signals using 488 nm laser into channel 2 (Ch02) through 528/35 bandpass filter. Speed beads (400041, Merck) were used for instrument alignment and were run constantly during image acquisition for focusing and camera alignment; Speed beads were visualized in channel 6 (Ch06). Channels 1 and 9 (Ch01 and Ch09) were used for bright field (BF) images and were used for camera alignment. Images were acquired using ISX.ex software and data were analyzed using IDEAS.exe (V 6.2) software. Using ISX, Ch02 intensities (log scale) of all events were displayed in a histogram. Speed beads bleach weak single up to the intensity of 1e4 in Ch02, therefore, intensities of 1e^4^ and higher were gated as GFP-positive signals, and 500-1000 GFP positive images were collected. Images were then imported to IDEAS where they were displayed as a function of Aspect ratio (y-axis) and Area (x-axis) in a dot plot. Singlets were gated between 0-50 X-axis and 0.87-1.0 Y-axis co-ordinates, while doublets were gated between 0-50 X-axis and 0.5-0.87 Y-axis co-ordinates. Events outside of these coordinates were considered as unresolved and were not included in data analysis. In contrast to the analysis presented on the *sar*A-GFP reporter strain, imaging flow cytometry analysis with USA300 LAC cells labelled with RSG was not conclusive, since the fluorescent signal was not strong enough to distinguish bacterial cells from autofluorescent beads present in the calibration fluid.

### Specific experimental procedures for analysis of growth heterogeneity in USA300 LAC colonies

#### Sample preparation

By default, single colonies of strain *S. aureus* USA300 LAC grown overnight (3 biological replicates) were suspended in PBS (D8537, Sigma) and mixed well using vortex mixer. Suspended bacteria were sonicated for 5 minutes at room temperature (RT) using Branson 3510 water sonicator to break bacterial clumps. 1e8 cells were stained with 0.1 mM *Bac*Light RedoxSensor Green (RSG, B34954, Invitrogen) for 10 minutes at 37°C in dark. For RSG titration, 1e^8^ cells were stained with 0.00, 0.05, 0.1, 0.2 and 0.5 μM of RSG, as above. The cells were diluted to 1e^7^ cell suspension in 1ml PBS, stained with 0.067 mM propidium iodide (PI, B34954, Invitrogen) for 5 minutes at RT, and sonicated for 1 minute prior to flow cytometric analysis. Unstained cells were used to set up flow cytometer gates and were processed as above.

For experiments to distinguish populations derived from different locations within colonies of 24 or 48 h of age, material was carefully harvested either from the center vs. edges of the colonies, or from the top vs. the bottom. These cells were prepared for flow cytometry analysis without RSG-labeling.

#### FACS-sorting and downstream analysis

The blue laser (488 nm) was used to collect Forward scatter (FSC) and Side scatter (SCC) through photodiode, and the RSG signal (FITC/GFP) through a 530/10 bandpass filter; The yellow-green laser (561 nm) was used to collect PI signals through a 582/15 filter. Threshold operator or triggering threshold was set to 200 on SSC. PMT voltages were adjusted for optimal separation of different populations as 650 (FSC), 420 (SSC), 650 (GFP), and 850 (PI). Single events were sorted into 96-well plates for growth analysis in liquid culture following the general description above.

### Specific experimental procedures for analysis of dormant growth variants elicited by low pH

#### Sample preparation

*S. aureus* individual colonies (3 biological replicates) were inoculated into the high glucose (4.5 g/L) DMEM media (with phenol red, L-glutamine, and sodium bicarbonate, without sodium pyruvate from Life Technologies) supplemented with 10% FBS. pH of this media was adjusted to pH 5.5 with citric acid. After incubation at 37 °C in 5% CO2 for 48 hrs samples were vortexed followed by centrifugation at 5000rpm for 5 min at room temp. Supernatants were discarded and bacterial pellets were washed with pre-warmed PBS and the cell concentration was adjusted to McFarland 0.5. Next, samples were labelled with BacLight™ RedoxSensor™ Green (Invitrogen™) at a final concentration of 200 nM and incubated in dark at 37°C for 10 min with moderate shaking. After 10 min, propidium iodide (PI) at final concentration 20 μM was added to the sample and incubated in the dark at 37°C for 10 minutes.

#### FACS-sorting and downstream analysis

Samples were then analyzed by BD FACSAria III instrument (BD Biosciences). Neutral density (ND) filter 1.0 was used and the channel voltages were set as follows: Forward scatter (FSC, 200 V), side scatter (SSC, 350 V), threshold (FSC/SSC, 200) and a flowrate 1.0 to analyze the bacterial cell size, aggregation complexity, and background noise. Cellular fluorescence for RedoxSensor™ Green reagent was collected with blue laser 488nm and 530/30 bandpass filter, whereas fluorescence from PI–stained samples was collected with a 561 nm excitation yellow-green laser and the (PE)-Texas Red (610/20 nm band-pass filter). A total of 50000 events were recorded for post-data analysis. BD FACSDiva software V 9.0 was used to gate the target population of interest and sorting. Prior to target population gating, background noise was excluded from data by gating only cell population using FSC and SSC. Then RSG-stained live population cells were distinguished from PI stained dead population by their distinct FITC values (x-axis) against PE-TexRed values (y-axis) at logarithmic scale. After assessment of viability, the live RSG^+^, PI^-^ populations were gated and prepared for index sorting with single cell precision mode with a 70 μm nozzle. Sorting was performed on single well agar plate and 96 well plate with pre-added TSA for growth dynamic analyses in both liquid and solid media following the general procedures described above. Any post-analysis and visual representation of flow cytometry data was performed with Flowjo software. Data from agar were counted for events in three categories: (a) Regular colony, (b) Small colony, and (c) No colony. For liquid growth analysis, the cell fates were (a) Regular growth, (b) Delayed growth, and (c) No growth.

### Specific experimental procedures for antibiotic dependent colony variant vancomycin-intermediate S. aureus strain (VISA)

#### Sample preparation

Single colonies (3 biological replicates) of the vancomycin susceptible *S. aureus* strain ATCC 29213 and representative clinical MRSA-VISA strain Mu50 (ATCC 700699) were grown overnight in Tryptic Soy Broth (TSB) at 37°C. Prior to labeling with Vancomycin-BODIPY FL (Vanco-BFL), both *S. aureus* strain were grown to exponential phase and cell concentration was adjusted to McFarland 0.5. Next, samples were labelled with Vanco-BFL at a final concentration of 1 μg/ml and incubated in the dark at 37°C for 30 min, before samples were additionally labelled with propidium iodide (PI) at final concentration 40 μM (adjusted for exponential phase) and incubated in the dark at 37°C for 10 minutes prior to flow cytometry analysis.

#### FACS-sorting and downstream analysis

Vanco-BFL signal was collected with blue laser 488nm and 530/30 bandpass filter, whereas fluorescence from PI–stained samples was collected with a 561 nm excitation yellow-green laser and the (PE)-Texas Red (610/20 nm band-pass filter). FSC and SSC was used to target the main cell events excluding the background noise. PI-negative cells were gated and sorted with index sorting (single cell precision mode activated from sorting layout in the BD FACSDiva software) onto Mueller Hinton agar (MHA) supplemented with different concentrations of vancomycin (0, 0.5, 2 or 8 μg/ml). Colony growth was evaluated after 24 and 48 h.

Cellular phenotypic profiling and backtracing analysis was performed with VISA cells sorted out onto MHA with 2 μg/ml vancomycin. After 48 h incubation at 37°C, the cell fates were assessed as (a) regular growth, (b) colony size variant, and (c) no growth phenotypes. Indexed phenotypes were further traced back in flow cytometry data for finding ancestral relation. For all sorted events in the different cell fate groups, the cellular Vanco-BFL fluorescence intensity values were retrieved for group analysis in Prism 9.5.

For assessment of growth kinetics of VISA strain Mu50 in the presence or absence of vancomycin, cells were sorted onto MHA or MHA + 2 μg/ml vancomycin and colony growth were monitored using time-lapse imaging data as described above.

## Author contributions

Conceptualization - C.S.L., Data collection - J.H., B.S. and C.A., Coding and testing of time-lapse image analysis program – T.H., A.A.S., Script for single cell-derived liquid culture analysis – A.M., Data analysis – all authors, Visualization- J.H., B.S. and C.S.L, supervision, funding acquisition - M.J., K.H. and C.S.L. Manuscript draft – J.H., B.S., C.S.L., Manuscript editing – All authors.

## Acknowledgements

We thank Dr. Alexander Horswill for pCM29-p*sarA*-GFP plasmid.This work was funded by the Norwegian Research Council (NFR, project 319829) and a Centre for New Antibacterial Strategies (CANS) starting-grant through the Trond-Mohn Foundation to C.S.L, and HNF1475-19 to M.J.

