## Supplementary Information for "High-throughput single-cell phenotypic profiling and backtracing exposes and predicts clinically relevant subpopulations in isogenic *Staphylococcus aureus* communities"

### Supplementary Figures

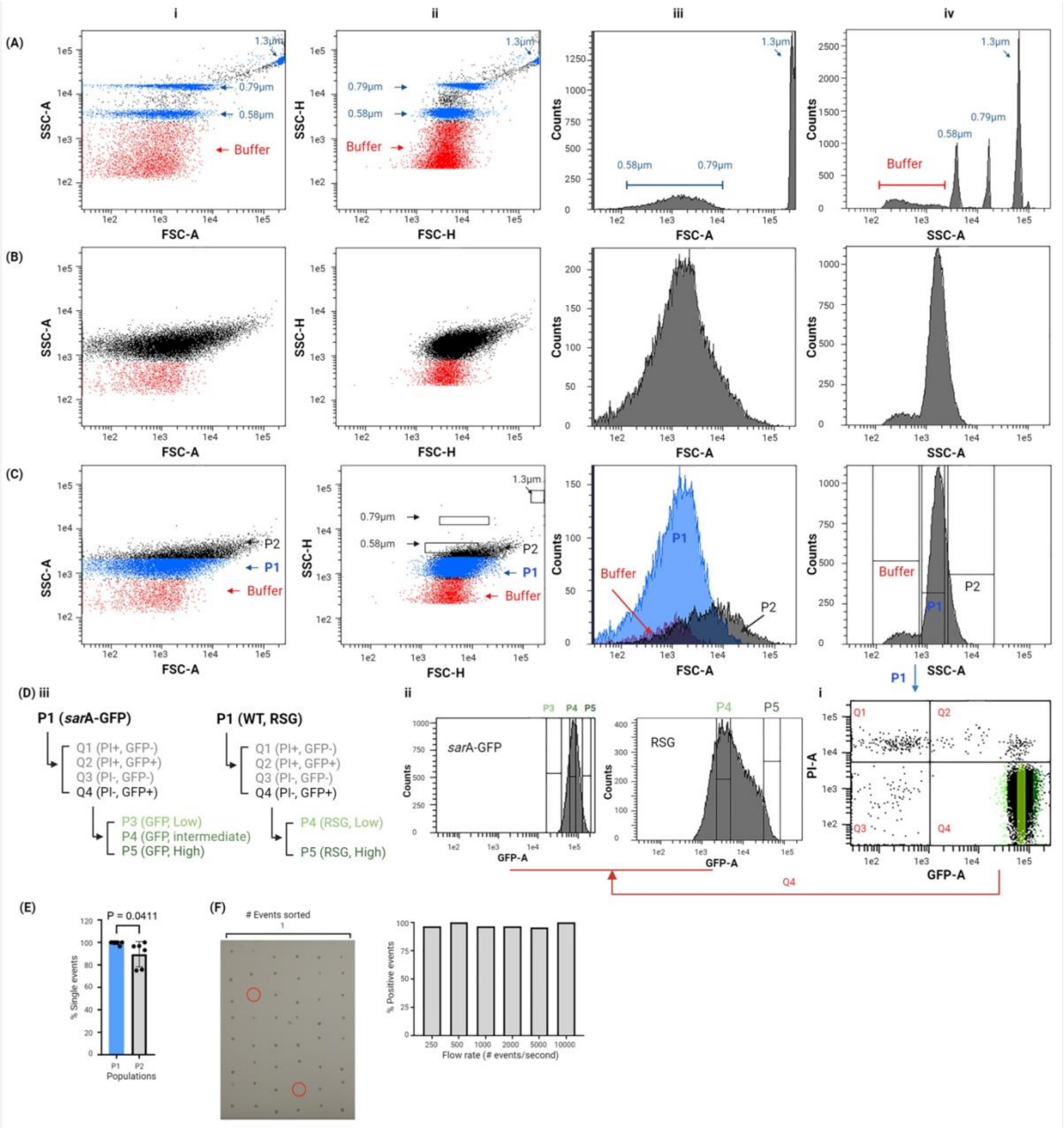


**Figure S1. Gating scheme and gate-hierarchy for experiments with *sar*A-GFP reporter strain**: (A) Size beads displayed as SSC-A and FSC-A (i), displayed as SSC-H and FSC-H (ii), FSC profile (iii) and SSC profile (iv). (B & C) comparison of bacteria displayed as SSC-A and FSC-A (i), displayed as SSC-H and FSC-H (ii), FSC profile (iii) and SSC profile (iv). Bacterial populations were gated as P1 and P2 on SSC-A plot (c-iv). Buffer is shown in red, P1 and P2 populations are shown in blue and black, respectively. (D) Downstream gating of P1 population for sorting: (D-i) P1 population displayed as PI-A and GFP-A to split P1 population as PI+GFP/RSG- (Q1), PI+GFP/RSG+ (Q2), PI-GFP/RSG- (Q3) and PI-GFP/RSG+ (Q4). (D-ii) GFP fluorescence of Q4 population was further displayed as GFP-A histogram and cell population was gated for their fluorescence intensity as low (GFP/RSG), intermediate (GFP/RSG) and high (GFP/RSG). (D-iii) overall gating hierarchy for *sar*A-reporter and WT cells. (E) Percent events showing single CFU events following single events sorting from P1 and P2 populations. (F) photograph of CFUs emerging on agar plates following single sorting events. Red circles mark sorted events not leading to colony growth. Accompanied graph shows percent of successful single CFUs emerged following single event sorting at different flow rates.

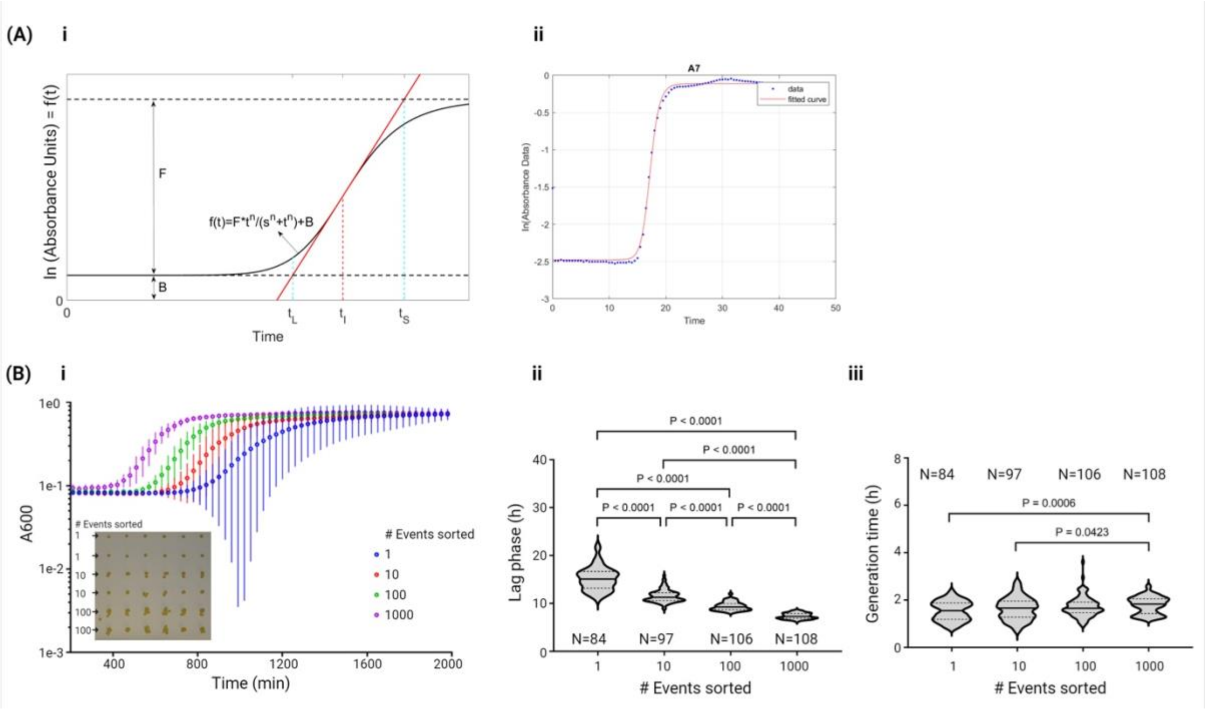


**Figure S2: Quantification of single cell-derived liquid growth cultures. (A) Fluorescence Non-Linear Regression Analysis.** (i) Based on fluorescence data expressed in log absorbance units (y-axis) versus time (x-axis), the duration of the lag phase ($t_L$), onset of the stationary phase ($t_S$) and the growth rate are defined in terms of a Hill function $f(t) = \frac{F*t^n}{(s^n+t^n)} + B$ where t is time, F the maximum fluorescence relative to baseline B and s is the timepoint at which the fluorescence level is at F/2 + B. The growth rate is defined as the slope of the tangent line of the Hill function at the inflection point, $t_I$, whereby $t_L$ and $t_S$ are defined as the intersection of the tangent line with the baseline B and the maximum fluorescence level (F+B), respectively. (ii) Illustration of the outcome of the non-linear regression to actual data. (B) Broth culture analysis of events sorted in multiples of 10. (i) Average A600, (ii) lag phase (h) and (iii) generation time (h) from 4 biological replicates. Inset: photograph of CFUs obtained from 1-100 sorted events following overnight growth.

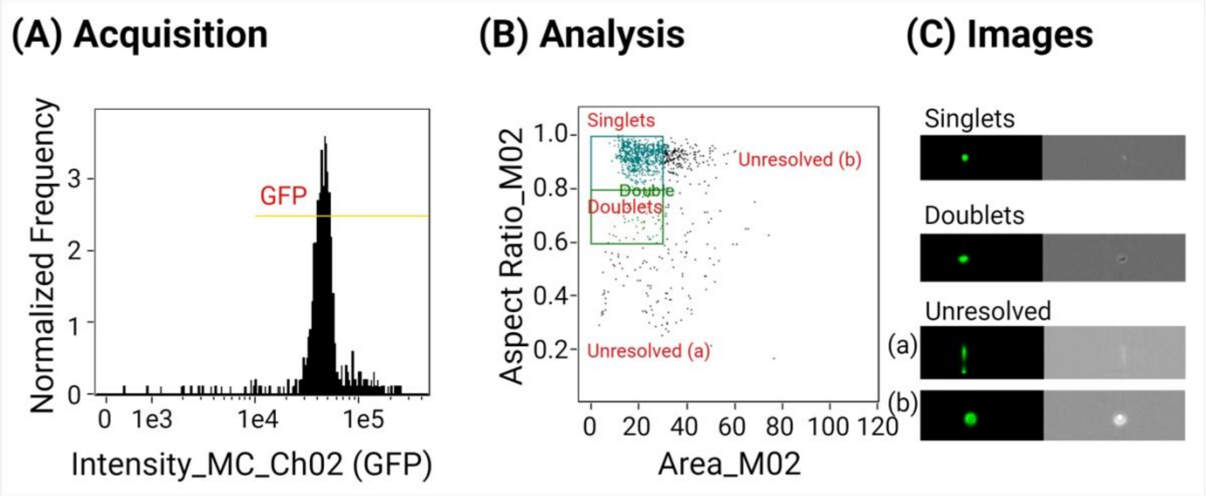


**Figure S3. ImageStream-analysis of FACS-purified cells**. (A) GFP intensity of all events displayed as normalized frequency and intensity of Ch02 (GFP channel). Events with GFP intensity higher than 1e$^4$ were gated as GFP+ and data from these events were acquired. (B) GFP+ events displayed as aspect ratio and area. Events were gated as singlets (co-ordinates, X-axis 0-50, Y-axis 0.87-1.0) and doublets (co-ordinates, X-axis 0-50, Y-axis 0.5-0.87). The remainder of the events was considered as unresolved and were not included in analysis. Events were considered unresolved when they either failed to form a circular shape (a) or to focus properly (b). (C) GFP (left) and bright field (right) images of selected cells from singlets, doublets, and unresolved populations.

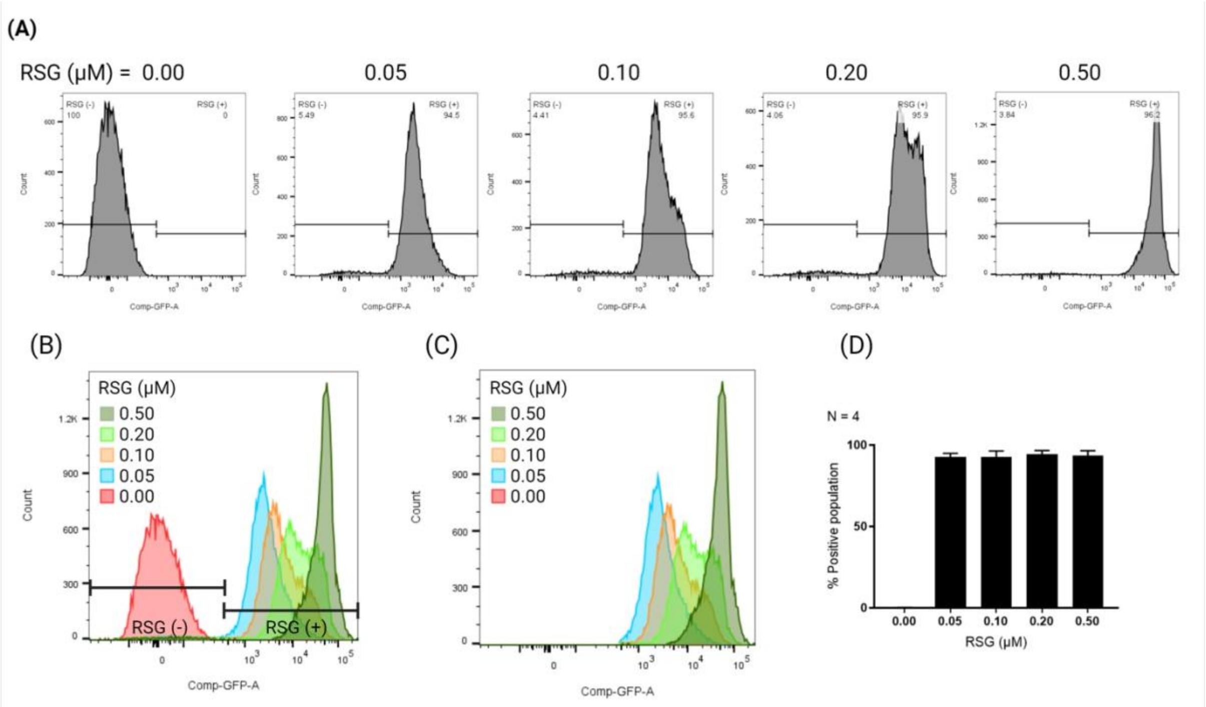


**Figure S4**. RSG titration. Flow cytometry histograms of a colony of *S. aureus* USA300 LAC profile stained in dose-response with different concentrations of RSG. (A) Individual histograms with RSG$^{(-)}$ and RSG$^{(+)}$ gating, (B) Overlay of histograms shown in A (C) Overlay of only RSG$^{(+)}$ population stained with different RSG concentrations. (D) Combined analysis of the RSG-positive population (N=4 biological replicates).

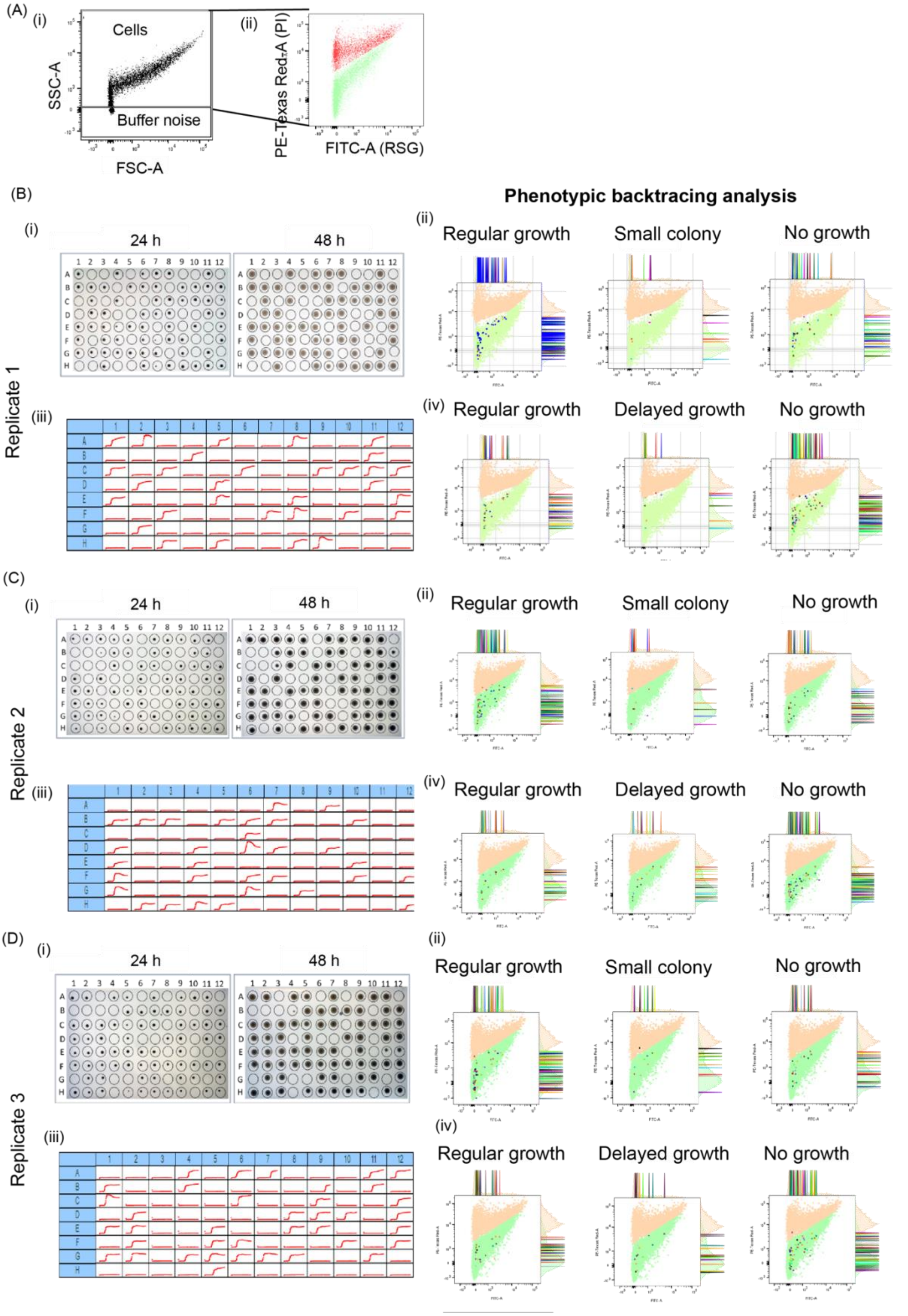


**Supplementary Figure S5. Low pH influence on *S. aureus* single cell growth phenotype on solid and in liquid media. (A)** Flow cytometry dot plot represents (i) Gating of all cellular event excluding background noise (ii) Gated population of RSG positive metabolic active population (green) and PI positive dead population (red). **(B-D)** An overview of the downstream growth analysis post sorting of single cells biological replicates 1 (B), 2 (C) and 3 (D). (i) Colony growth on agar plate after 24h and 48h . (ii) Phenotypic backtracing analysis of agar growth analysis. The graph shows a flow cytometry dot plot (x-axis: RSG, y-axis: PI) highlighting the location of the events of the different cell fate categories (Regular growth, small colony, no growth) on the background of the general cell population ($PI^-$: green, $PI^+$: orange). iii) Single-cell derived growth curves over 48h in TSB in a 96-well plate read-out. (iv) Phenotypic backtracing analysis of single-cell derived growth in liquid culture. The graph shows a flow cytometry dot plot (x-axis: RSG, y-axis: PI) highlighting the location of the events of the different cell fate categories (Regular growth, delayed growth, no growth) on the background of the general cell population ($PI^-$: green, $PI^+$: orange)

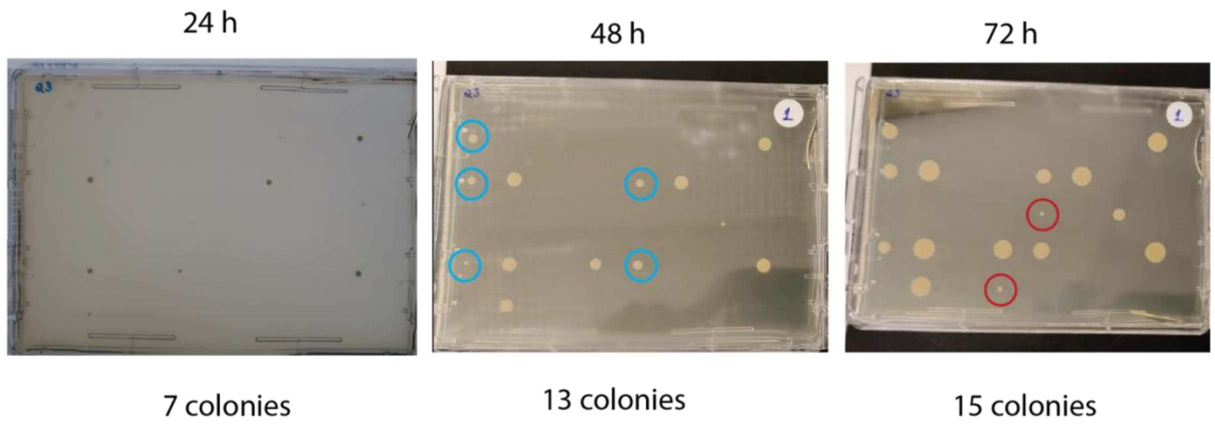


**Figure S6. Examples of *S. aureus* colonies appearing after 48 h.** Exemplary images of *S. aureus* colonies appearing after prolonged incubation times. *S. aureus* LAC was grown in DMEM with 10% FBS media adjusted to pH 5.5 for 72 h, before 48 $PI^-$-negative cells were sorted onto a rectangular TSB plate for cultivation at 37 °C. Colony growth was monitored at the indicated time points. The coloured circles indicated colonies appearing for the first time at 48 h (blue) and 72 h (red), respectively.

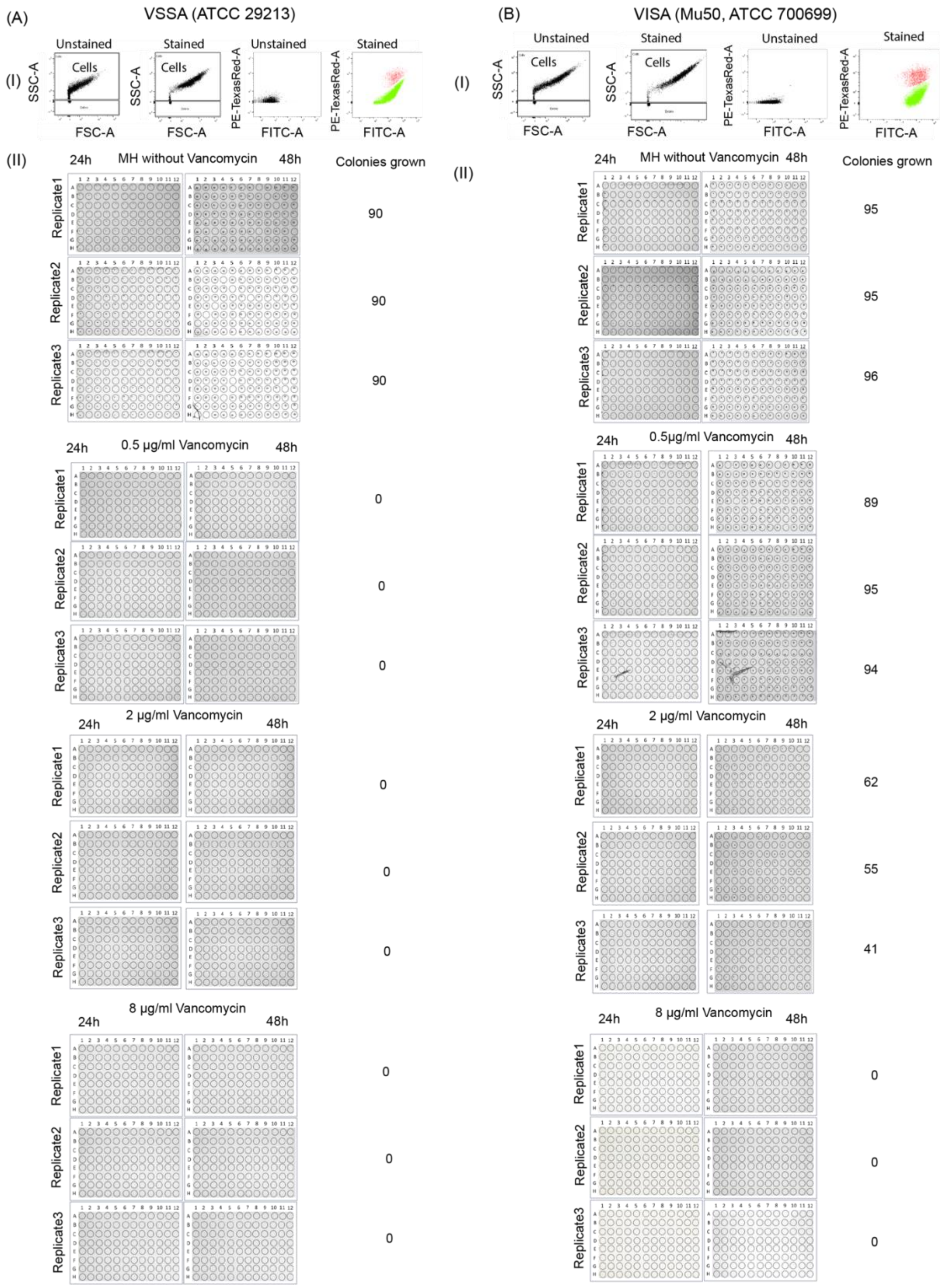


**Supplementary Figure S7. Cellular phenotypic profiling coupled to single-cell derived vancomycin susceptibility testing.** (A) VSSA strain ATCC 29213 and (B) VISA Mu50 ATCC 700699 cells were labelled with Vanco-BFL and propidium iodide for flow cytometry analysis and single-cell sorting for growth analysis in the presence of different concentrations of vancomycin (0, 0.5, 2 or 8 μg/mL). (i) Flow cytometry dot plot represents gating of all cellular events under stained or unstained condition, and Vanco-BFL positive (blue) and PI positive (putatively dead) population (red). (ii) Colony growth analysis after sorting of single cells onto MHA plates supplemented with the indicated different concentrations.

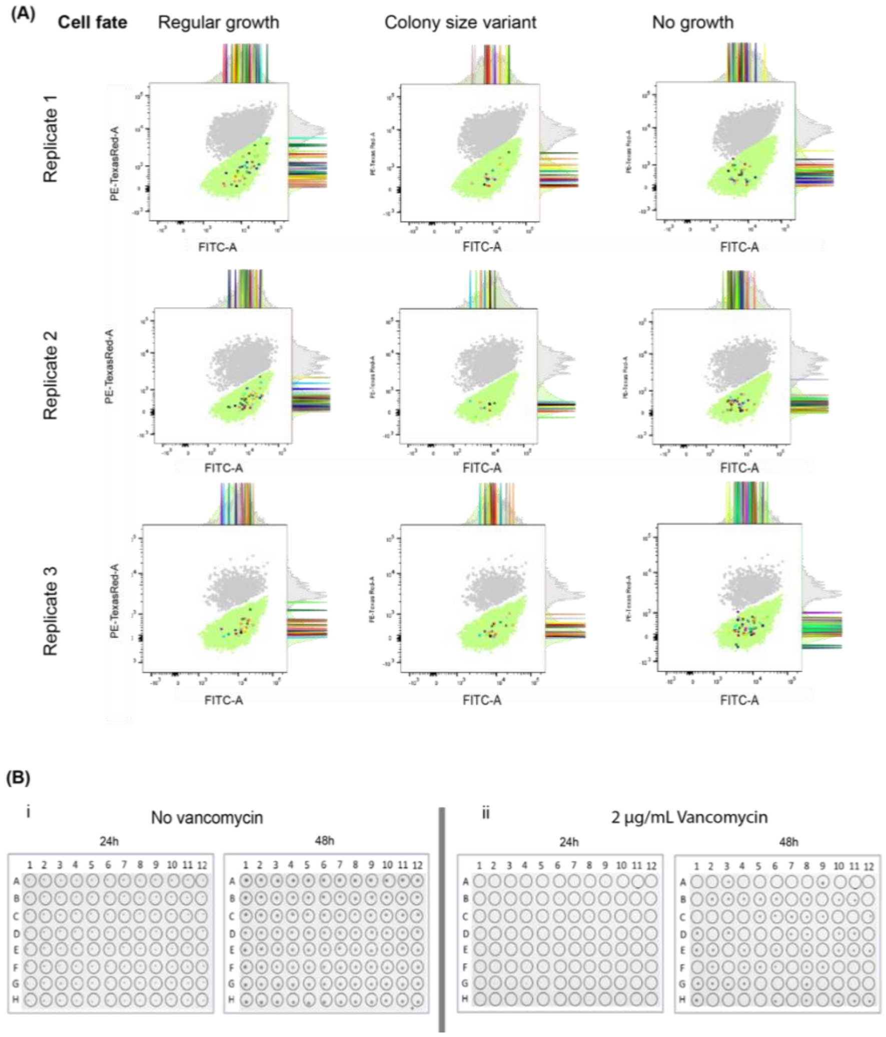


**Supplementary Figure S8. Phenotypic backtracing analysis of cells with different susceptibility to vancomycin.** VISA Mu50 ATCC 700699 cells were labelled with Bodipy-BFL and PI and single $PI^{(-)}$ cells sorted onto MH agar plates with or without 2 µg/ml vancomycin for 24 or 48h to determine cell fates (Regular growth, colony size variant, no growth). (A) Flow cytometry dot plots of cells highlighting the location of the events of the different cell fate categories on the background of the general cell population (x-axis: Vanco-BFL, y-axis: PI, background colors: $PI^-$ - green, $PI^+$ - red). A total N=96 different events each are shown for 2 biological replicates. (B) Single frame image extracted from time-lapse video of agar plates depicting colony growth of Vanco-BFL sorted VISA Mu50 on (i) MH agar plate without and (ii) with vancomycin.
